# Liquid chalk is an antiseptic against SARS-CoV-2 and influenza A respiratory viruses

**DOI:** 10.1101/2020.11.02.364661

**Authors:** Julie L. McAuley, Joshua M. Deerain, William Hammersla, Turgut E. Aktepe, Damian J.F. Purcell, Jason M. Mackenzie

## Abstract

The COVID-19 pandemic has impacted and enforced significant restrictions within our societies, including the attendance of the public and professional athletes in gyms. Liquid chalk is a commonly used accessories in gyms and is comprised of magnesium carbonate and alcohol that quickly evaporates on the hands to leave a layer of dry chalk. We investigated whether liquid chalk is an antiseptic against highly pathogenic human viruses including, SARS-CoV-2, influenza virus and noroviruses. Chalk was applied before or after virus inoculum and recovery of infectious virus was determined to mimic the use in the gym. We observed that addition of chalk before or after virus contact lead to a significant reduction on recovery of infectious SARS-CoV-2 and influenza but had little impact on norovirus. These observations suggest that the use and application of liquid chalk can be an effective and suitable antiseptic for major sporting events, such as the Olympic Games.

## Introduction

The use and application of hand sanitizers in preventing the spread of infectious microbial diseases is primarily based on the 60-80% v/v of alcohol included within these products [1, 2]. The use of hand sanitizers has been an important control measure in limiting the spread of virus during the recent COVID-19 pandemic in some settings [3, 4]. Liquid Chalk is a product that comprises of magnesium carbonate (chalk), 40-80% alcohol (generally ethanol, methanol or isopropanol), water and sometimes other additives including resins or proprietary materials, depending on the manufacturer. When applied to the hands the liquid chalk is distributed across the surface of the hand and then dries into a thin chalk layer as the alcohol evaporates. Due to the high percentage of alcohol, liquid chalk has been suggested to act as a hand sanitizer against SARS coronavirus, although this has yet to be proven experimentally. In this study we investigated and evaluated the application of various liquid chalk products as antiseptics against the spread and transmission of SARS-CoV-2 (the causative agent of the COVID-19 pandemic), influenza A virus (H1N1) (IAV) and norovirus, using the surrogate model of mouse norovirus (MNV).

## Materials and Methods

### Cell culture maintenance and virus stocks

Vero cells (American Type Culture Collection [ATCC]) were maintained in Minimal Essential Media (MEM) supplemented with 10% heat-inactivated foetal bovine serum (FBS), 10 μM HEPES, 2 mM glutamine and antibiotics ((100 units/mL Penicillin G, 100 μg/mL Streptomycin). Madin-Darby canine kidney (MDCK) cells were grown in Roswell Park Memorial Institute (RPMI) media supplemented with 10% FBS, 2 mM glutamine and antibiotics. RAW 264.7 cells were maintained in Dulbecco’s Modified Eagle’s Medium (DMEM) with 10% FBS and 1% GlutaMAX. Cell cultures were maintained at 37°C in a 5% CO_2_ incubator. SARS-CoV-2 isolate hCoV-19/Australia/VIC01/2020 [5] stocks, were produced as previously described [6]. The influenza A virus isolate, A/Puerto Rico/8/34 was produced as previously described [7]. Details of MNV strain CW1 have been described previously [8, 9].

### Cytotoxicity assay 96® Non-Radioactive Cytotoxicity Assay (Promega)

Vero, MDCK and RAW cells were plated in a 96-well plate to 80% confluency. 50ul of various liquid chalk samples were aseptically air-dried, resuspended in 500ul of cell culture media and centrifuged at 400xg for 3 mins to remove excess chalk particles (this sample is depicted as “neat”). Neat supernatant was 10-fold serially diluted in respective tissue culture media, added to the 96-well plate containing cells and incubated at 37°C for 24 hours. Following the incubation period, 10ul of 10x Lysis Solution was added for 45 mins to the control wells to generate a Maximum LDH Release Control. 50ul of supernatant from each sample (in duplicates) was transferred to a fresh 96-well flat clear bottom plate and incubated for 30 mins with 50ul of CytoTox 96® Reagent. To end the reaction, 50ul of Stop Solution was added to each well and absorbance was recorded at 490nm.

Percent cytotoxicity = 100 × (Experimental LDH Release (OD490)/Maximum LDH Release (OD490))

### Chalk exposure assays

All SARS-CoV-2 infection cultures were conducted within the High Containment Facilities in a PC3 laboratory at the Doherty Institute. For the chalk first assay, sample of the liquid chalk was aseptically smeared onto 2-4 discrete areas covering approximately 2cm round surface (approx. 50ul) on a sterile tissue culture dish and allowed to dry. 50uL virus inoculum was then applied. Where the inoculum did not absorb into the dry chalk, a slurry of chalk:virus mixture was created using a sterile tip and mixing. 15min later, 500uL infection media (identical to culture media but without the presence of sera) was added then mixed with the chalk:virus sample and collected. Excess chalk was pelleted at 400 g for 3min, then a 50% tissue culture infectious dose (TCID_50_) assay performed on the supernatant as previously described [6, 7]. For the virus first assay, 50uL virus inoculum was added to 2-4 discrete areas on a sterile tissue culture dish, then chalk added and spread to cover an approximate 2cm round surface on a sterile tissue culture dish. After 15min incubation at room temperature, 500uL infection media was added and sample mixed, then remaining virus present in the sample quantitated in the same way as per the chalk first procedure. For both tests, no chalk control received 50uL infection media.

### Statistical Analyses

Data is representative of at least 2 independent experiments and was analysed using GraphPad Prism v8.0.

### Patient and Public Involvement statemen

We wish to state that patients or the public were not involved in the design, or conduct, or reporting, or dissemination plans of our research

### Ethical approval information

No animals or humans were utilised during this study. The infection with SARS-CoV-2 is covered by our Institutional Biosafety Committee approval number 2020/028.

## Results

### Experimental rationale

In our experimental design we investigated two different approaches to represent the scenarios encountered within the gym environment. Firstly, we tested the application of liquid chalk on a surface that already contained the virus, and secondly, we tested the ability of liquid chalk to prevent transmission once applied to a surface with subsequent addition of the virus. For the virus first experiments, a volume containing a known titre of virus (SARS-CoV-2, IAV or MNV) was applied to a plastic surface. Subsequently, a known volume of four commercially available and widely used liquid chalk products was added, smeared to thin layer and then allowed to dry. The entire layer was then resuspended in tissue culture media and the chalk particulate was removed by centrifugation. The resultant supernatant was then diluted and added to Vero (SARS-CoV-2), MDCK (IAV) or RAW264.7 cells (MNV) and virus infectivity was measured by TCID_50_/mL. For the chalk first experiments, the exact same procedure described above was performed except the volume of chalk was applied first and allowed to dry and then the volume of virus was added to the dried chalk.

### Liquid chalk prevents the recovery of infectious SARS-CoV-2

As can be observed in Figure 1, all four liquid chalk products significantly impacted the recovery of SARS-CoV-2 virus from the surface to which it was applied. Intriguingly, Chalk #1-3 all displayed complete loss of the recovery of virus (to the limit of detection) whereas Chalk #4 also had a significant impact, but some residual virus could be recovered. Of interest was our observation that the application of liquid chalk before or after virus inoculum had an equal effect on the recovery of SARS-CoV-2 virus. In this assay, the alcohol contained in the liquid chalk product was allowed to evaporate prior to contact with virus, suggesting the dried chalk provided a viricidal activity. To confirm the antiviral effects of alcohols, we treated SARS-CoV-2 virus preparations with differing amounts and types of alcohols (including those commonly found in Liquid Chalk products). As summarised in Table 1, we observed that all alcohols had a virucidal effect on SARS-CoV-2. Thus, our results indicate that the application and implementation of Liquid Chalk can be a suitable antiseptic against the transmission of SARS-CoV-2. Liquid chalk itself was not cytotoxic to any of the cell types used in this study at the concentrations shown to be effective in reducing virus recovery (Supplementary Figure 1).

**Table 1.**
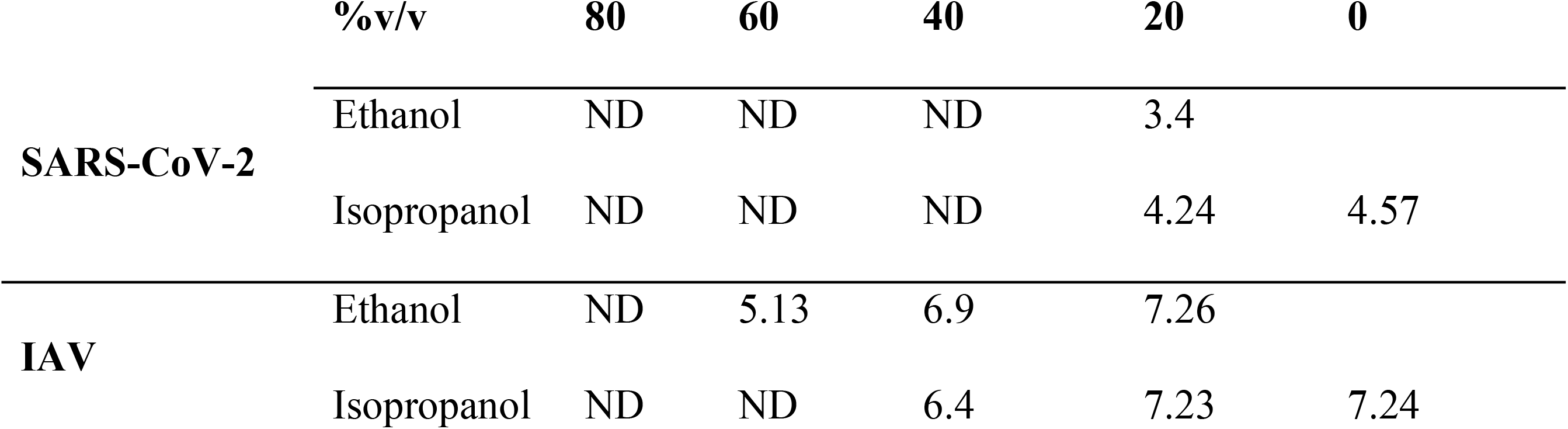
Recovery of virus (Log_10_/mL) after 15min exposure to alcohol.

**Figure 1:**
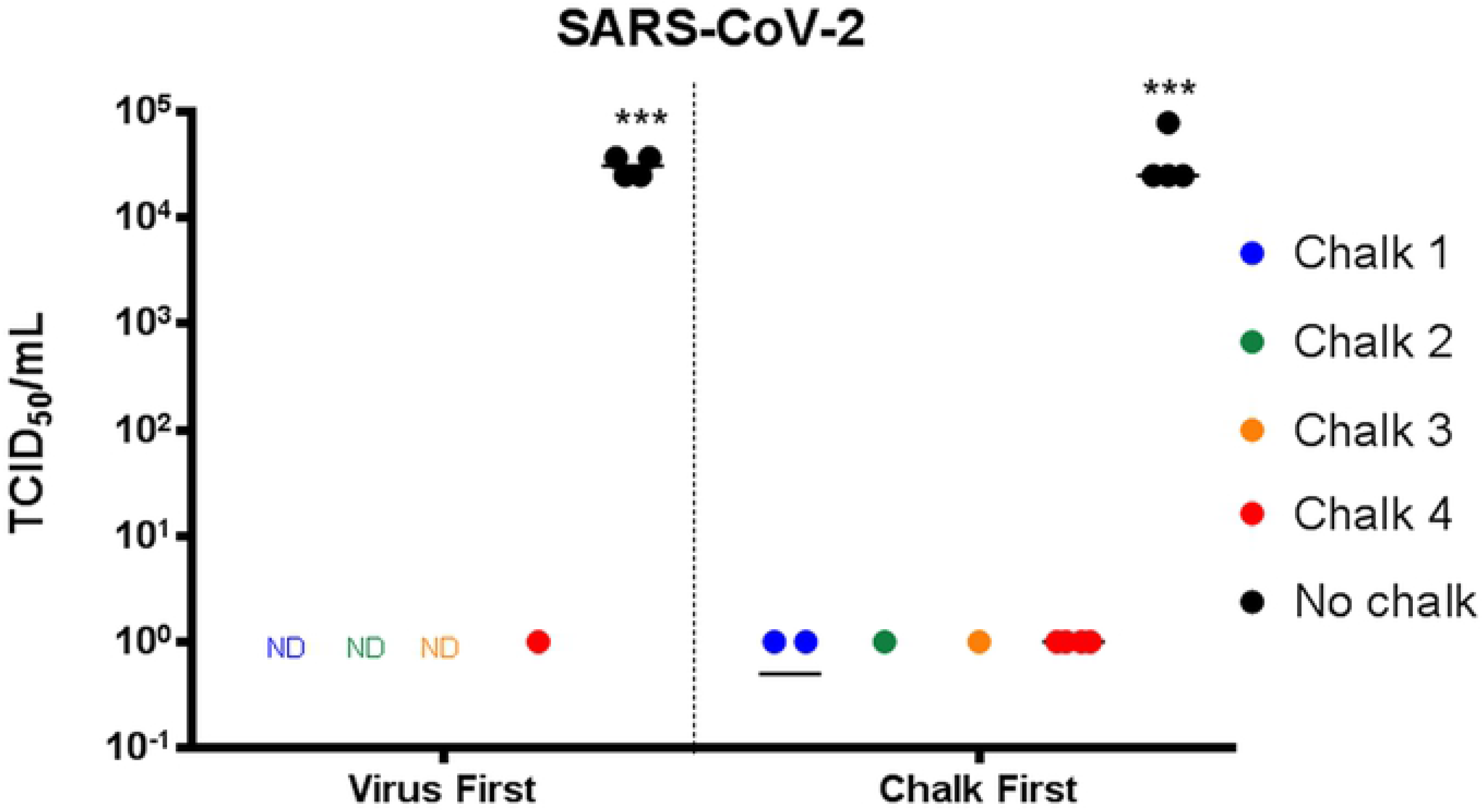
SARS-CoV-2 is rendered non-infectious by gym Liquid Chalk. All chalks tested significantly reduced the amount of infectious SARS-CoV-2 in the sample compared to the no chalk control (ND = not detectable) when added either before or after the viral inoculum. (*** *p* < 0.001 compared to no chalk, one-way ANOVA).

### Liquid chalk prevents the recovery of infectious influenza virus (IAV)

To further our study, we also tested the antiviral effect of Liquid Chalk against another highly infectious and pathogenic respiratory viral pathogen IAV. The experiments were performed exactly as described above and the virus TCID_50_/mL was titrated on MDCK cells. As can be observed in Figure 2, all four Liquid Chalk products were effective in restricting the recovery of IAV compared to SARS-CoV-2. However, for IAV the effect was greater when the chalk was applied to the virus inoculum rather than chalk first then virus, linking the presence of alcohol as a significant antiseptic component against IAV. As can be observed application Chalk #1 and 3 reduced the virus recovery by approximately 6 logs (TCID_50_/mL) whereas Chalk #2 and #4 reduced the recovery by ~2.5 logs. When chalk was first applied to the surface, all chalks induced an approximate 2-4 log decrease in virus recovery. Overall, these results indicate that the application of liquid both to IAV or onto a surface can reduce the recovery of IAV.

**Figure 2:**
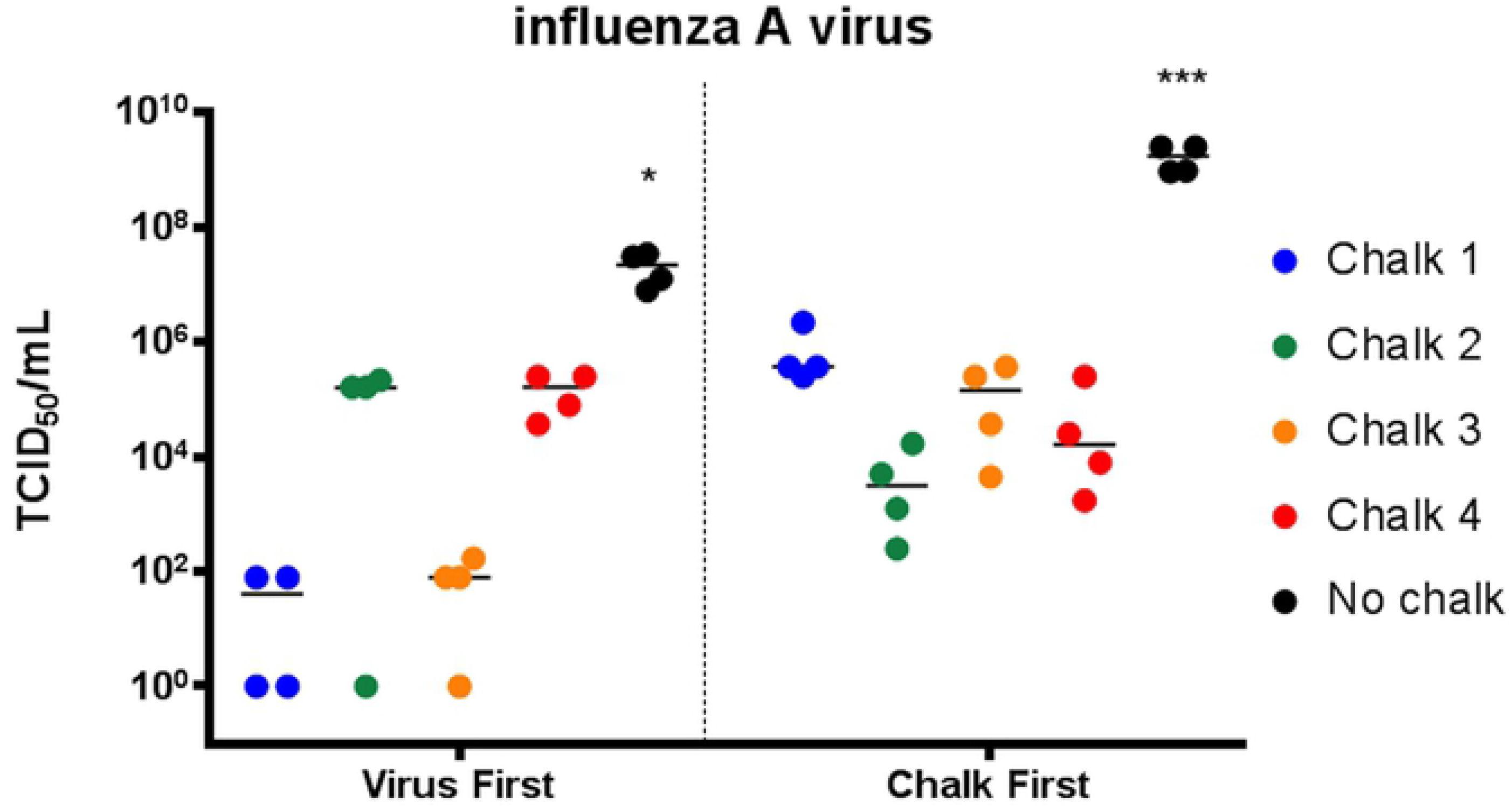
Influenza A virus is significantly inactivated by Liquid Chalk. When influenza A virus was applied to the surface first, chalk 1 and chalk 3 reduced the amount of infectious virus to nearly undetectable levels (*p* < 0.001 compared to no chalk control). When chalk was applied first and dried, the recovery of infectious influenza A virus was significantly reduced compared to the chalk control, but markedly more infectious virus remained in the samples treated with chalk 1 and chalk 3 compared to the virus first samples that underwent the same treatment (data not significant). (\**p* < 0.05, *** *p* < 0.001 compared to no chalk, one-way ANOVA).

### Liquid chalk does not prevent the recovery of infectious norovirus

As a comparator, we also investigated the ability of Liquid Chalk to inactivate another highly infectious viral pathogen, norovirus. As human norovirus is difficult to cultivate in laboratory conditions, we utilised the widely appreciated surrogate murine norovirus (MNV) for our studies [10]. Again, the experiments were identical to those described above except the viral TCID_50_/mL was performed on RAW264.7 (murine macrophage) cells. Intriguingly, Figure 3 shows that MNV is relatively resistant to the virucidal properties of the Liquid Chalk products. We observed that Chalks #1-3 had very little impact on virus recovery, with a 0.5 log reduction the best that we observed. However, in contrast to SARS-CoV-2 we observed an approximate 1 log reduction upon application of Chalk #4. This is interesting as the major difference between MNV and SARS-CoV-2 and IAV is that MNV is a non-enveloped virus, whereas the other two contain a host-derived lipid membrane as their outer most layer. Thus, it would be interesting to identify the constituents of Chalk #4 as it was the effective agent against MNV and but was observed to be slightly less effective against SARS-CoV-2.

**Figure 3:**
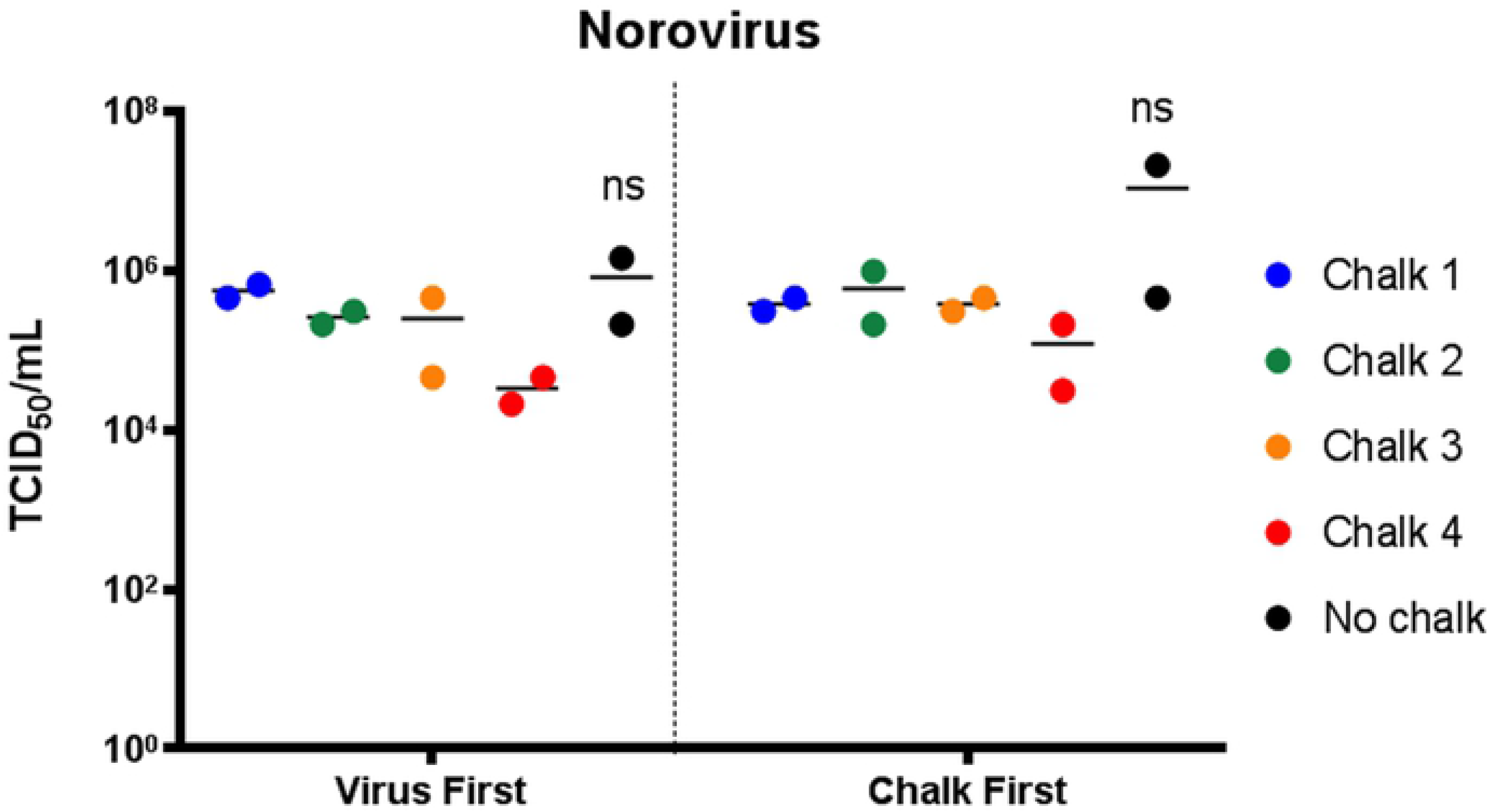
Norovirus remains infectious when exposed to Liquid Chalk. Norovirus was not rendered non-infectious when treated with gym chalk, regardless of whether the virus was added to dry chalk, or chalk was added to virus inoculum (*p* > 0.05 to no chalk, one-way ANOVA).

### SARS-CoV-2 and IAV are sensitive to treatment with various concentrations of alcohols

As different alcohols are the major constituents of liquid chalk, we additionally evaluated the impact of alcohol alone on the recovery of infectious virus. As can be observed in Table 1, exposure of SARS-CoV-2 and IAV to both ethanol and isopropanol is detrimental to the infectiousness of these viruses up to a dilution of 40% v/v for IAV and 20% v/v for SARS-CoV2. Given proprietary information regarding the alcohol content of chalks, we can only conjecture that if lower than these percentages of alcohol are present in the Liquid Chalk, then it is likely that the antiviral activity we observe can be attributed to the chalk component.

## Discussion

There have recently been two press releases, one using seasonal CoV and not SARS-CoV-2 [11], while the other used SARS-CoV-2 but only a single chalk product [12]. Neither public release investigated the efficiency against other highly infectious viral pathogens. To our knowledge this study is the first to demonstrate antiviral activity of a range of commercially available and commonly used Liquid Chalks. Given the uncertainty of re-opening gyms due to contact transmission from potentially contaminated equipment, our findings that Liquid Chalks have anti-viral activity against SARS-CoV-2 may aid in decision making for re-opening gyms in the future. This is important due to the impact of gym closures (due to COVID-19), on personal fitness, professional sports and particularly mental health and well-being.

## Acknowledgements

The authors acknowledge the contribution and assistance of Melbourne Health through its Victorian Infectious Diseases Reference Laboratory at the Doherty Institute, in providing our laboratory with isolated SARS-CoV-2 material.

## Author Contributions

JLM, TEA, JMD, DFJP and JMM designed the experimental plan; JLM, TEA and JMD performed then experiments themselves; WH provided the liquid chalk; JLM, TEA, JMD, and JMM collated and analysed the data and wrote the manuscript

## Funding

The University of Melbourne acknowledges the support of a grant administered by the State Government of Victoria to JMM

## Conflicts of Interest

The authors declare no conflict of interest. The funders had no role in the design of the study; in the collection, analyses, or interpretation of data; in the writing of the manuscript, or in the decision to publish the results.

## Footnotes

### Competing interests

All authors declare they have no competing interests with this study

### Data sharing

All results gained during this study will be made available via journal access after acceptance and publication of the article on open access university website, or can be obtained by the senior author.

This information has not been presented at any meetings to date

## References

1. Golin AP, Choi D, Ghahary A. Hand sanitizers: A review of ingredients, mechanisms of action, modes of delivery, and efficacy against coronaviruses. American journal of infection control. 2020;48(9):1062–7. Epub 2020/06/18. doi: 10.1016/j.ajic.2020.06.182. PubMed PMID: 32565272.

2. Kampf G. Efficacy of ethanol against viruses in hand disinfection. Journal of Hospital Infection. 2018;98(4):331–8. doi: 10.1016/j.jhin.2017.08.025.

3. Kratzel A, Todt D, V’kovski P, Steiner S, Gultom M, Thao TTN, et al. Inactivation of Severe Acute Respiratory Syndrome Coronavirus 2 by WHO-Recommended Hand Rub Formulations and Alcohols. Emerging Infectious Disease journal. 2020;26(7):1592. doi: 10.3201/eid2607.200915.

4. Leslie RA, Zhou SS, Macinga DR. Inactivation of SARS-CoV-2 by commercially available alcohol-based hand sanitizers. American Journal of Infection Control. 2020. doi: https://doi.org/10.1016/j.ajic.2020.08.020.

5. Caly L, Druce J, Roberts J, Bond K, Tran T, Kostecki R, et al. Isolation and rapid sharing of the 2019 novel coronavirus (SARS-CoV-2) from the first patient diagnosed with COVID-19 in Australia. Med J Aust. 2020;212(10):459–62. Epub 2020/04/03. doi: 10.5694/mja2.50569. PubMed PMID: 32237278; PubMed Central PMCID: PMCPMC7228321.

6. Lee JYH, Best N, McAuley J, Porter JL, Seemann T, Schultz MB, et al. Validation of a single-step, single-tube reverse transcription loop-mediated isothermal amplification assay for rapid detection of SARS-CoV-2 RNA. J Med Microbiol. 2020. doi: 10.1099/jmm.0.001238. PubMed PMID: 32755529.

7. McAuley JL, Hornung F, Boyd KL, Smith AM, McKeon R, Bennink J, et al. Expression of the 1918 influenza A virus PB1-F2 enhances the pathogenesis of viral and secondary bacterial pneumonia. Cell Host Microbe. 2007;2(4):240–9. Epub 2007/11/17. doi: 10.1016/j.chom.2007.09.001. PubMed PMID: 18005742; PubMed Central PMCID: PMCPMC2083255.

8. Karst SM, Wobus CE, Lay M, Davidson J, Virgin HW, IV. STAT1-Dependent Innate Immunity to a Norwalk-Like Virus. Science. 2003;299(5612):1575–8.

9. Hyde JL, Sosnovtsev SV, Green KY, Wobus C, Virgin HW, Mackenzie JM. Mouse norovirus replication is associated with virus-induced vesicle clusters originating from membranes derived from the secretory pathway. J Virol. 2009;83(19):9709–19. Epub 2009/07/10. doi: 10.1128/JVI.00600-09. PubMed PMID: 19587041; PubMed Central PMCID: PMCPMC2748037.

10. Wobus CE, Thackray LB, Virgin HWt. Murine norovirus: a model system to study norovirus biology and pathogenesis. J Virol. 2006;80(11):5104–12. PubMed PMID: 16698991.

11. Research Shows Chalk could be a climbers best Friend [Internet]. https://www.abcwalls.co.uk/wp-content/uploads/Corona-Virus-and-Chalk-Press-release-v2.pdf; 2020

12. Casey C. Liquid Chalk Proven in CU Labs to Kill Coronavirus, Potentially Helping Climbing Gyms to Safely Reopen https://news.cuanschutz.edu/news-stories/liquid-chalk-proven-in-cu-lab-to-kill-coronavirus-potentially-helping-gyms-to-safely-reopen?utm_source=miragenews&utm_medium=miragenews&utm_campaign=news2020.

